# Festa: *F*Lexible *E*Xon-Based *S*Plicing And *T*Ranscription *A*Nnotation

**DOI:** 10.1101/300947

**Authors:** Rago Alfredo, Cobourne John K.

**Affiliations:** Department of Biology, Lund University, Lund, Sweden; School of Biosciences, University of Birmingham, Birmingham, UK

## Abstract

We introduce FESTA, an R based algorithm that allows detection of alternative splicing based on experiment-specific exon expression data. FESTA disentangles alternative splicing signal from whole-gene transcription, facilitating the discovery and characterization of novel regulatory events even in the absence of transcript annotations or paired-end data. We also include customization options to increase its applicability on different platforms and experimental designs as well as a tool for the conversion from transcript expression to inclusion ratios.

## 1 Background

Alternative splicing is a widespread feature of eukaryotic gene regulation which can be represented as a two step process. Transcription generates the total amount pre-mRNA per gene locus and splicing determines the proportions of each alternative transcript that is produced. Based on this model we can distinguish between two types of exons. **Constitutive exons** are present in all transcripts produced from a gene Chen, 2013 and their expression levels match the total production of transcripts from that gene. **Facultative exons** are instead present in only some transcripts, and their expression is instead determined by the joint action of RNA production and splicing.

Commonly used transcript quantification methods discard information contained in constitutive exons or average it to match the proportions provided by transcript-specific exons (Trapnell *et al.*, 2010). These practices have two main drawbacks. First, quantifying transcript-specific expression from alternative exons effectively reports the joint effect of transcription and splicing dependent signal, confounding different biological processes. Second, if reads are mapped to pre-annotated transcripts novel transcriptional events might be missed entirely. Dataset-specific estimation of constitutive and transcript-specific exons is therefore advisable for the discovery of novel alternative splicing events relevant to the design of interest (i.e. Dai *et al.*, 2012; McManus *et al.*, 2014).

Correlation based exon clustering provides a simple method for dataset-specific estimation of new splicing events (Patrick *et al.*, 2013). Since each exon’s expression is proportional to the sum of the expression of each of the transcripts which include it, exons which share transcripts will have stronger correlations that exons which don’t. The exon clusters obtained via correlation clustering will each represent either a group of exons which characterizes a specific transcript or the group of constitutive exons, which is included in all transcripts. Since constitutive exon clusters are present in all transcripts they will have higher (or tied) expression values compared to all other exon clusters, allowing their identification. Despite being intuitive and effective, the results of correlation-based hierarchical clustering depend on the tree-cut height, which determines how different two exons need to be in order to be assigned to different groups. However, there is so far no guide criterion for determining an appropriate threshold.

In this paper, we present a simple algorithm that solves this issue by setting gene-specific thresholds based on highly customizable biological expectations. We also provide a function to calculate inclusion ratios of alternative exon groups in order to allow analysis of transcription independent effects of splicing.

## 2 Implementation and Usage

### 2.1 Data input and filtering

FESTA requires two input files: an exon by sample expression table and an exon to gene assignment table. In order to avoid spurious grouping resulting from correlations in the noise component we advise thresholding raw expression data, removing all values that score below minimal signal and excluding all exons that lack expression in a sufficient number of biological replicates for at least one of the dataset’s conditions.

### 2.2 Isoform detection

Figure 1 shows an outline of the FESTA algorithm, which is applied to iteratively to each gene. If a gene has more than one expressed exon, FESTA calculates a clustering tree based on the correlation matrix of expressed exons. FESTA then cuts the tree at the lowest level (one exon per cluster) and ranks each group’s expression in each sample (figure 1, A).

**Figure 1:**
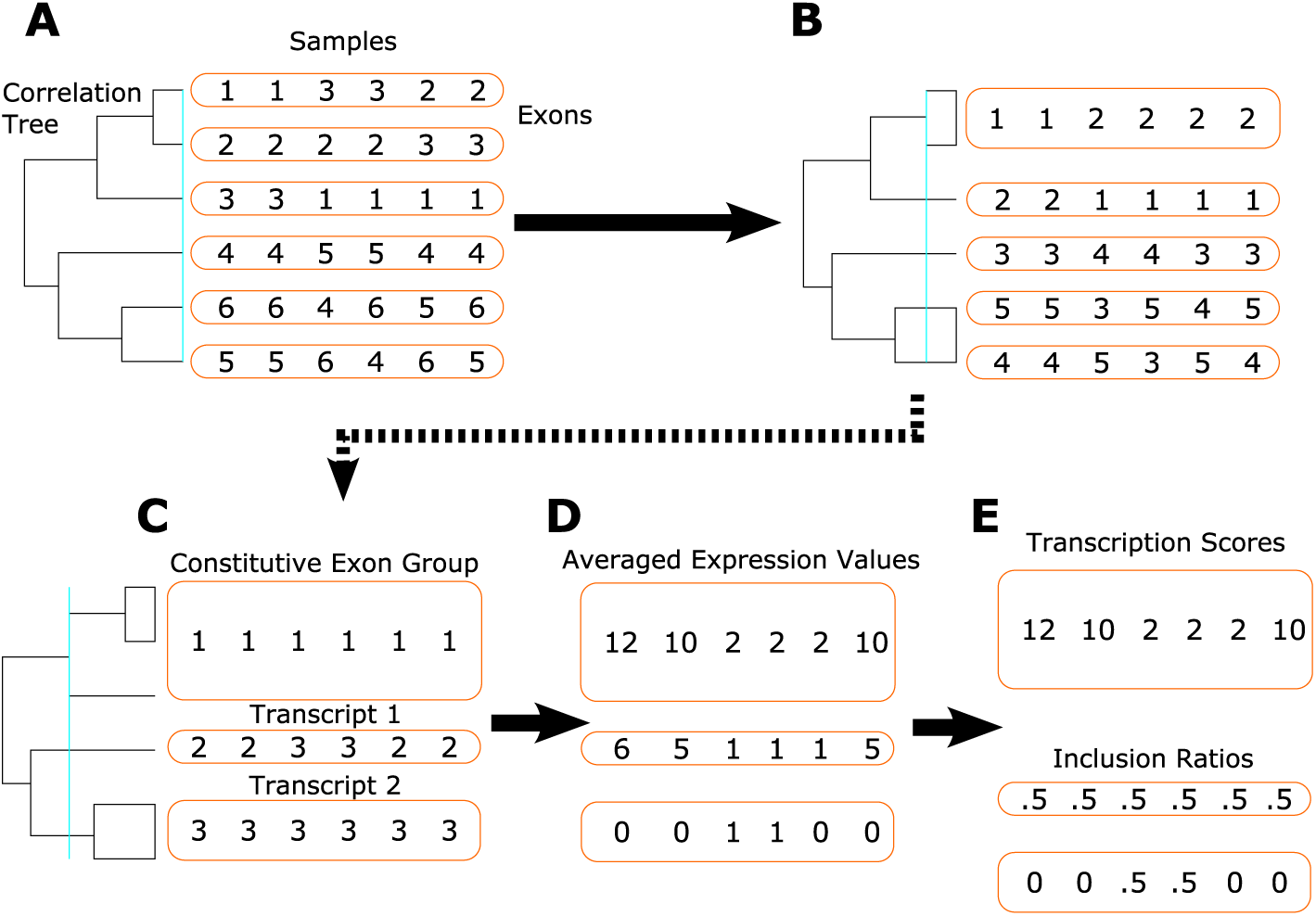
Outline of the FESTA algorithm. Steps A-C are handled by the ClusterExons function. Step D and optional step E are handled by the AverageExons function.

If any exon group is ranked first or tied for first across all samples we consider it to be the constitutive part of the gene, record the cluster assignments at the tree cut level and proceed on to the next gene. If no exon group ranks as first or tied across all samples, FESTA moves up a level in the hierarchical clustering tree, averages expression scores in exon groups with more than one exon and re-calculates the exon group rankings across the dataset (figure 1, B).

FESTA iteratively calculates the expression rankings of exon groups at each level until a single exon group shows the highest expression across all samples (figure 1, C). If no exon group meets constitutive exon criteria at the highest level, the algorithm converges on single-group clustering: all exons are annotated as constitutive and the gene is reported as lacking significant splicing events.

FESTA generates a single expression score for each group by averaging the expression scores of all its exons (figure 1, D). These raw expression scores can be directly used for analyses on individual transcript abundance. Alternative splicing events can also be converted to inclusion ratios by dividing them by the transcription score of their gene (figure 1, E). Inclusion ratios range between zero (if the isoform is absent) and one (if all transcripts produced by the gene include those exons) and can be used to analyze the effects of alternative splicing independently of the main gene’s overall expression.

### 2.3 Fine tuning parameters

Festa allows tuning of two main parameters which affect the sensitivity and power of the main clustering algorithm: significant digits and number of exceptions.

Significant digits allows the user to define numerical accuracy of expression measurements. Setting a high number of significant digits will result in less ties between exon groups but might cause over sensitivity to minor fluctuations in expression values between biologically co-expressed exons. Fewer significant digits increase the number of ties in rankings, decreasing the ability to differentiate constitutive exons from highly expressed alternative exons. The choice of this parameter should be based on an empirical estimation of the effective sensitivity of expression measurements in the dataset.

Number of exceptions allows to increase the permissiveness of constitutive exon group definitions. If this number is greater than zero, constitutive exons groups are allowed to be less expressed than other exons in a corresponding number of samples. For instance, setting exception number to 1 in a dataset with 25 samples will identify as constitutive any group of exons that is the most expressed (or tied) in at least 24 out of 25 samples; setting exception number to 2 will instead require only 23 out of 25 samples and so on. This parameter enables setting tree-cut height based on experimental design considerations, with higher values resulting in more isoforms and smaller constitutive exon groups.

### 2.4 Caveats

There are three caveats regarding FESTA’s current implementation. Firstly, the algorithm depends on the number of biological replicates to generate accurate exon rankings. Secondly, it does not currently make use of paired-end data. Lastly, as the algorithm attempts to identify isoform-specific exon groups it will not be able to detect isoforms characterized by different combinations of the same exons such as in the case of hypervariable combinatorial genes.

## 3 Conclusions

We present an intuitive method for the detection of transcription and splicing in transcriptomic data which requires only an exon by sample expression table. FESTA allows the end user to customize sensitivity using easily interpreted parameters which can be tuned to the experimental design and the instrument’s sensitivity. FESTA’s output is a reduced transcript by sample table, which retains only the splice variants observed in the experiments and can be directly-used in downstream analyses. The optional conversion from transcript abundances to splicing ratios allows the investigation of the effects of increasing the proportions of specific isoforms rather than their absolute abundances, allowing for a comparative study of the impact of transcriptional and splicing regulation.

## 4 Implementation

The package FESTA is available for R version 3 or over at: https://github.com/alfredorago/FESTA.

